# Low ultimate tensile strain in atherosclerotic plaques is linked to the presence of neovascularisation: insights from *ex vivo* uniaxial tensile testing

**DOI:** 10.1101/2025.06.18.660485

**Authors:** Y. Guendouz, F. Digeronimo, B. Tornifoglio, C. Lally

## Abstract

**Background:** Neovascularisation, the formation of new vessels within atherosclerotic plaques, has been associated with plaque progression and intraplaque haemorrhage. However, the relationship between neovascularisation and mechanical failure of plaques remains unclear. A better understanding of this link could improve rupture risk prediction beyond traditional reliance on arterial stenosis.

**Methods:** Human carotid plaques (n=7) from endarterectomy were sectioned into circumferential strips (n=25) and tested under uniaxial tension with digital image correlation (DIC) to determine ultimate tensile (UT) stress, UT strain, and elastic moduli. Immunohistochemistry with CD31 staining quantified neovascularisation. K-means clustering based on UT stress and strain identified mechanical phenotypes. A decision tree classifier was trained using neovascularisation percentage as the predictor.

**Results:** Neovascularisation showed a moderate negative correlation with UT strain (r = -0.46, p = 0.028), indicating plaques with greater neovascularisation failed at lower strains. K-means clustering identified two groups: Cluster 1 (lower UT strain) had significantly higher neovascularisation (median 2.21%) and greater initial stiffness than Cluster 2 (median 1.27%, p < 0.01). The decision tree using neovascularisation alone achieved 78.3% classification accuracy, with an area under the curve of 0.78. DIC analysis revealed rupture consistently occurred at regions of elevated local strain.

**Conclusion:** This study demonstrates a link between neovascularisation and mechanical failure in atherosclerotic plaques. Higher neovascularisation was associated with lower UT strain, suggesting it contributes to mechanical instability. These findings support the role of neovascularisation as a marker of plaque vulnerability and encourage its integration into ultrasound-based risk assessment strategies, such as contrast-enhanced ultrasound.

## 1. Introduction

Atherosclerosis remains one of the leading causes of cardiovascular morbidity and mortality worldwide^1^. Characterised by the build-up of plaques within the arterial walls, which can rupture. This rupture can cause the release of plaque material into the bloodstream, where they travel to smaller cerebral arteries, blocking blood flow and ultimately causing a stroke. Despite advances in cardiovascular care, predicting plaque rupture remains challenging, emphasising the need for better risk indicators beyond traditional metrics^2^.

Current clinical practice primarily assesses plaque vulnerability based on the degree of stenosis, the narrowing of the artery. However, research has shown that stenosis alone is insufficient to capture the complex behaviour of plaques and their rupture potential^2,3^. This underscores the need for a more comprehensive risk assessment of plaque rupture^4^.

In this context, several studies in the literature have turned their attention to exploring the microstructural characteristics of atherosclerotic plaques and the link to mechanical behaviour. For example, Johnston et al. used small angle light scattering to evaluate the dominant fibre orientation in plaque caps and found that caps with circumferentially aligned fibres were associated with greater strength^5^. Such findings support the need for further investigation into the local tissue microstructure to better understand its mechanical behaviour and its risk of failure^6–8^.

However, non-invasive imaging with the power to provide microstructural information which is indicative of mechanical stability and translatable to the clinic remains scarce. Maher et al. used clinical US imaging to classify plaques as either calcified, mixed or echolucent revealing that calcified plaques exhibited the stiffest behaviour while echolucent plaques were the least stiff^7^. Tornifoglio et al. used a specific MRI sequence, namely diffusion tensor imaging, to non-invasively obtain microstructural and mechanical insights. Although this technique is promising, translation from the lab to the clinic still requires considerable advancements, particularly in reducing the sequence acquisition time^8^.

Despite extensive efforts in the literature, a direct link between composition and plaque vulnerability remains lacking. As no current clinical imaging technique can directly correlate composition with the mechanical rupture risk of plaque, it is crucial to investigate the *ex vivo* mechanical characterisation of plaque incorporating microstructural information that could be translated into clinical practice.

Neovascularisation is defined as the formation of new microvessels within tissues, typically in response to stimuli such as hypoxia or inflammation^9^. Specifically, neovascularisation within atherosclerotic plaques represents a response to the increased metabolic needs of the progressing plaque, primarily driven by hypoxia^10^. In healthy arteries, oxygen and nutrients are delivered to the intima and media from the luminal side, while the adventitia is served by microvessels originating from the primary artery or neighbouring vessels^11^. Additionally, a decrease in arterial elasticity due to ageing or metabolic conditions reduces the natural flow driven by arterial pulsations. This flow aids nutrient transport across the vessel wall; however, its reduction further limits oxygenation, aggravating the hypoxic environment^11^. In atherosclerotic lesions, this situation intensifies, as expanding plaques not only restrict diffusion but also elevate local oxygen demand due to inflammation. The resulting hypoxia, particularly within the plaque core and adjacent media, promotes the release of pro-angiogenic factors like vascular endothelial growth factor, which drive the formation of fragile and permeable neovessels from the adventitia. Marsch et al. showed that hypoxia within plaques contributes to necrotic core expansion by impairing the process where macrophages clear apoptotic cells^12^. Their findings suggest that hypoxia drives plaque vulnerability highlighting neovascularisation as a potential marker of this process.

Studies also found that intraplaque haemorrhage (IPH) is prevalent in symptomatic carotid plaques, even in cases of mild stenosis, suggesting that factors beyond lumen narrowing can influence plaque vulnerability^13^. Neovascularisation plays a crucial role in this context, as the fragile microvessels formed within the plaque are prone to rupture, leading to IPH. The hypoxic environment stimulates neovascularisation which in turn can create a cycle that drives plaque vulnerability. Therefore, imaging of plaque neovascularisation may provide additional information about the presence of vulnerable plaques in patients.

US imaging, particularly contrast enhanced ultrasound (CEUS), has gained traction in clinical settings for visualising plaque neovascularisation^14^. CEUS enables detailed visualisation of intraplaque neovascularisation, offering a new dimension in characterising plaque composition and associated risks. The CEUS principle is based on micro-bubble contrast agents injected intravenously which enhance US signals by increasing blood backscatter^15,16^. Virmani et al. observed that angiogenesis can be the cause of IPH due to neovessel leakage and neovascularisation is also associated with plaque enlargement^17^.

CEUS also serves as a complementary modality alongside techniques like shear wave elastography (SWE) and has potential to be integrated in clinical risk assessments to help guide interventions aimed at preventing cerebrovascular events^18,19^. Furthermore, research has demonstrated a strong correlation between neovascularisation identified through histological analysis and CEUS findings^14,20^.

Although CEUS holds promise as a non-invasive method for assessing neovascularisation, robust evidence linking neovascularisation to plaque rupture risk remains limited^21,22^. This gap highlights the need for further research to support or enhance US-based risk assessment.

This study aims to bridge this gap by investigating the relationship between neovascularisation and plaque vulnerability. By employing uniaxial tensile testing, this study directly assesses plaque mechanical strength and failure while incorporating histological quantification of neovascularisation. This combined approach seeks to establish whether the presence and extent of neovascularisation can serve as an indicator of mechanical vulnerability, ultimately aiming to surpass the limitations of the measured degree of stenosis as a sole predictor of plaque rupture risk.

## 2. Materials and methods

An overview of the methods used in this study are shown in Figure 1.

**Figure 1:**
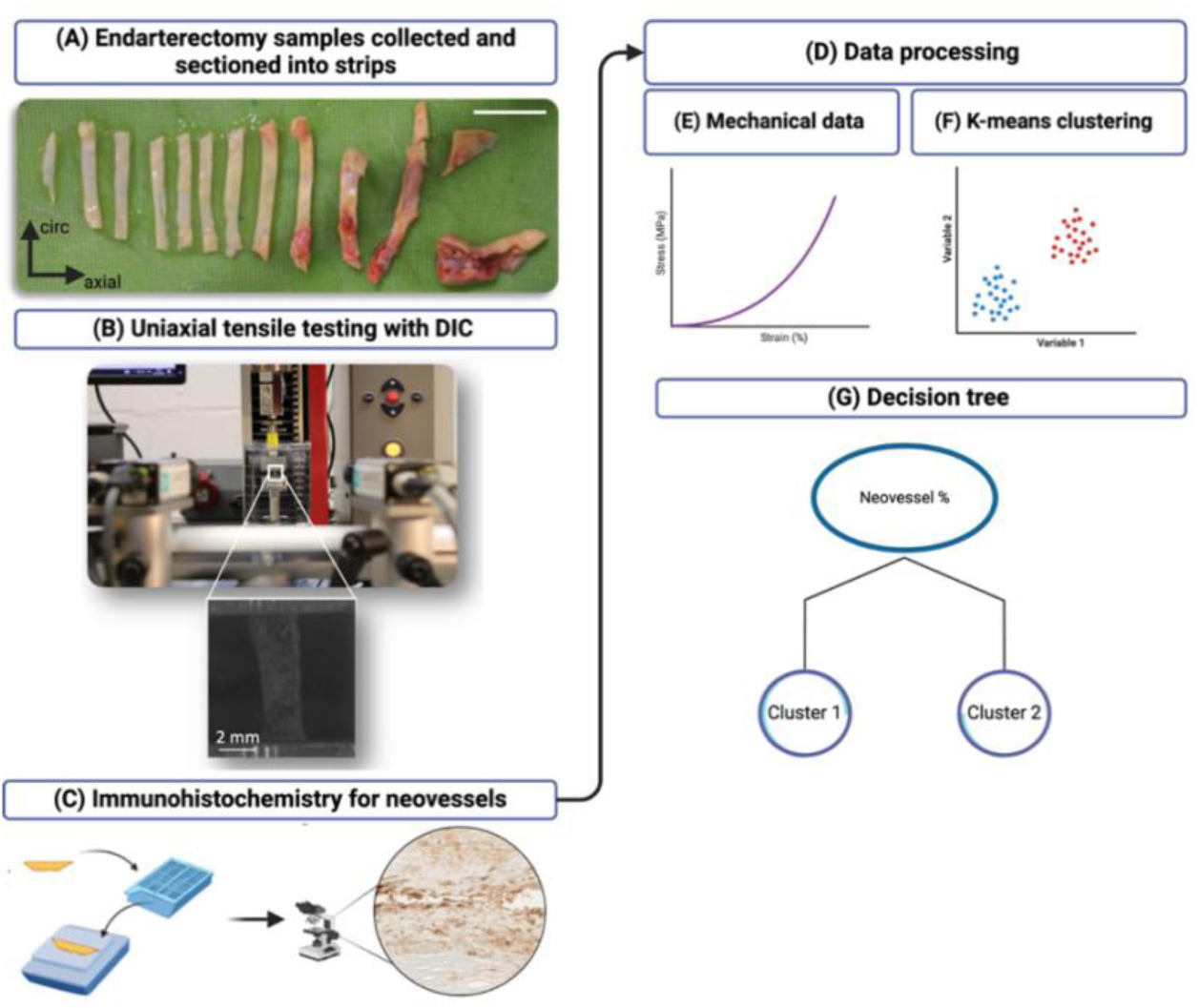
A) Excised endarterectomy carotid sample cut into strips. Scale bar: 1 cm. B) Photograph of strip mounted for uniaxial tensile testing. C) Schematic of immunohistochemistry workflow. D) Data processing flow. E) Representative stress-strain extracted from mechanical testing. F) K-means clustering. G) Decision tree classifier implemented to investigate neovascularisation as a classifier between the clusters.

### a. Sample acquisition

Carotid atherosclerotic plaques (n=7) were obtained from symptomatic carotid endarterectomy patients at the Galway Clinic, see Figure 2. Ethical approval was granted in accordance with the Declaration of Helsinki (2016-12 List 47 (4)). Plaques were rinsed in phosphate buffered saline (PBS), cryopreserved, and stored at -80°C until *ex vivo* testing.

**Figure 2:**
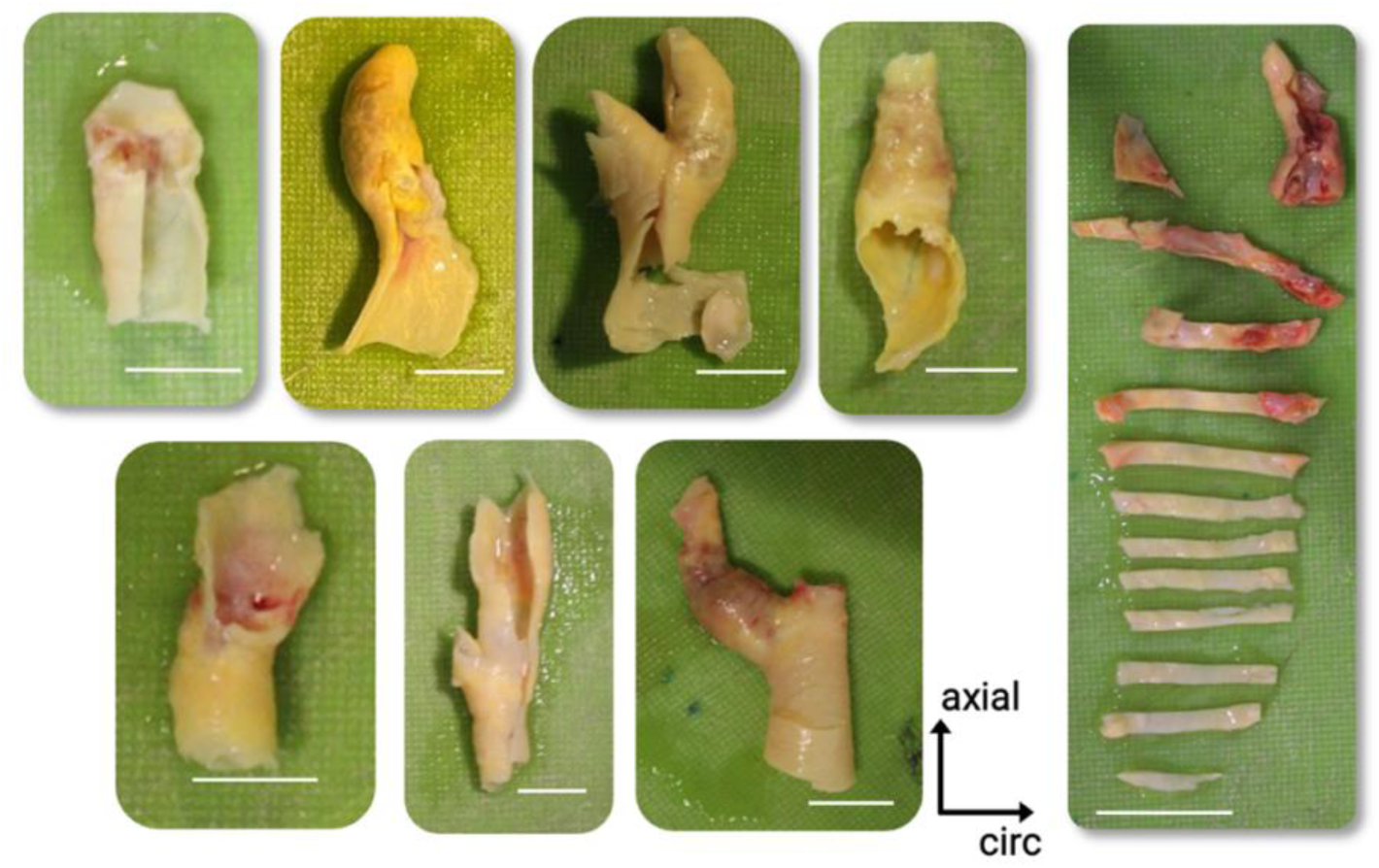
Photographs of atherosclerotic carotid plaques cut into strips for testing. Scale bars: 1 cm.

### b. Mechanical testing

#### i. Circumferential strips

Samples were thawed at 37°C and rinsed in PBS. Circumferential strips (n=38) were sectioned from the plaques, see Figure 3(1). Due to plaque tissue heterogeneity, dog-bone shapes - which are designed to minimise edge stress concentrations and ensure failure occurs at the centre of the gauge length - were not feasible. Plaque samples were cut into rectangular strips 2 mm in width, see Figure 3(2). The mean thickness for each strip was determined from three measurements taken with the Mitutoyo Litematic VL-50B for accuracy, allowing cross-sectional areas to be calculated. Of the 38 strips tested, 13 were excluded due to failure near the grips, issues with digital image correlation (DIC) analysis, or due to challenges in histological processing. To enable DIC analysis, the strips were sprayed with a tissue marking dye (Epredia™, Fisher Scientific) using an airbrush (Kkmoon Airbrush) connected to a single-cylinder piston compressor (ABEST). The workflow is illustrated in Figure 3.

**Figure 3:**
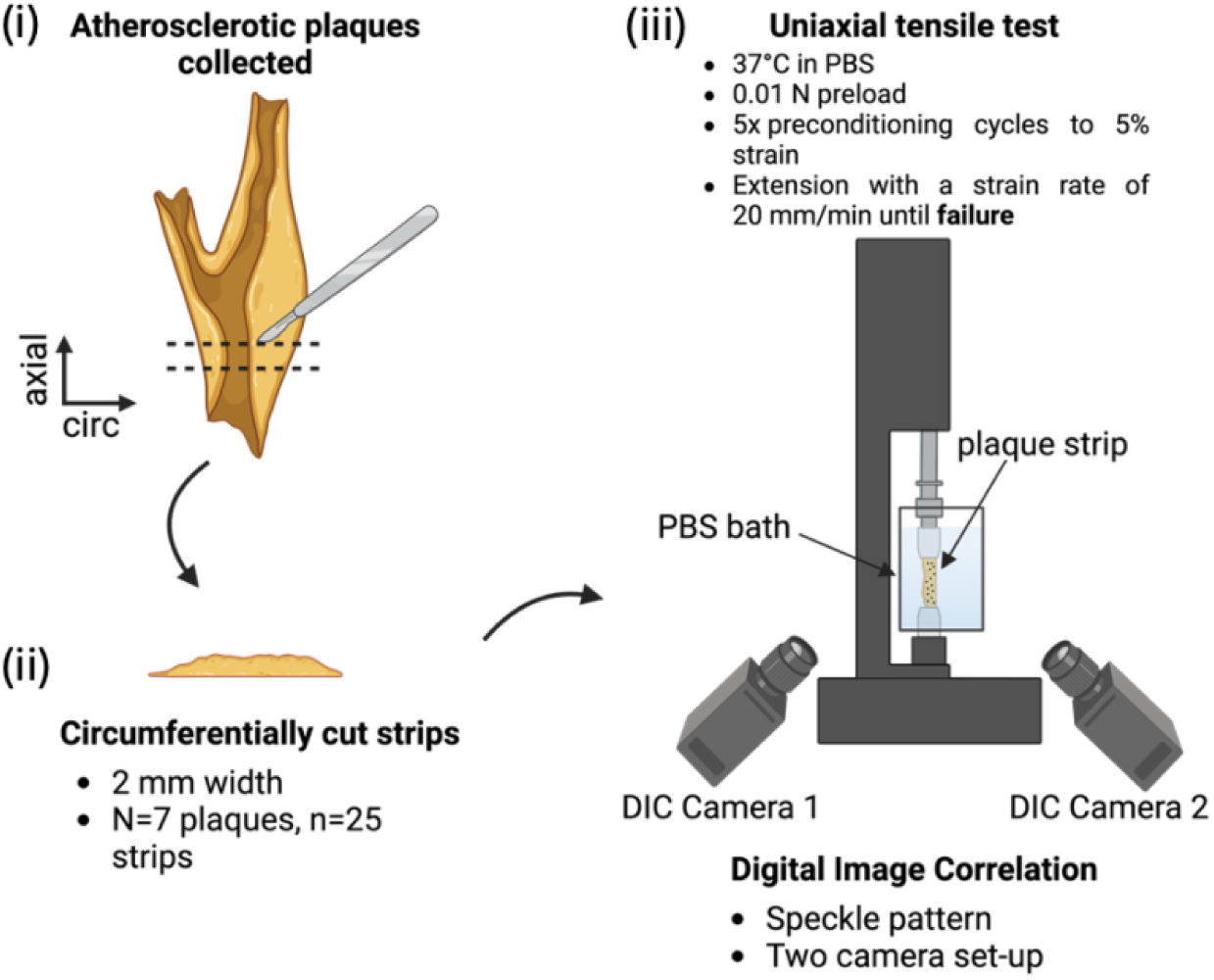
Framework for plaque testing.

#### ii. Uniaxial tensile test and DIC

Strips were subjected to uniaxial extension until failure using a uniaxial testing machine (Zwick Z005, Zwick GmbH & Co., Ulm, Germany). Strips were loaded in the circumferential direction to provide insights into the mechanical behaviour in the primary load-bearing orientation, see Figure 3(1,2). All tests were conducted in a PBS bath maintained at 37°C. The protocol started with a preload of 0.01 N, after which the force was zeroed, followed by five preconditioning cycles to 5% strain, and extension to failure, see Figure 3(3). Each step was performed at a strain rate of 20 mm/min. DIC was employed to track local strain deformations, using a two-camera system (Dantec Dynamics GmbH, Denmark) with prior calibration, see Figure 4. Images were captured at 5 Hz. After failure, all samples were fixed in 10% formalin for subsequent histological processing for no more than 48 hours.

**Figure 4:**
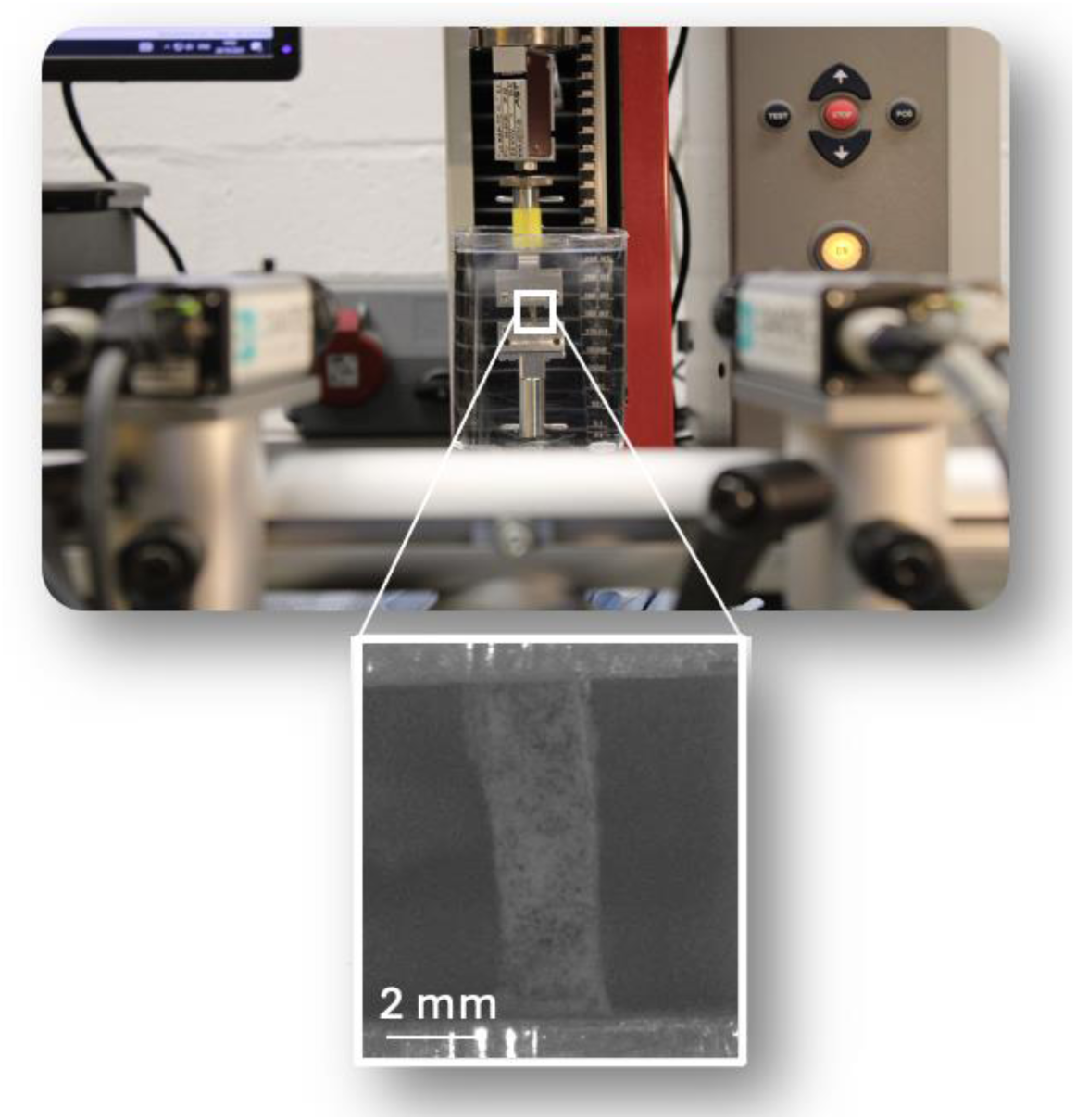
Experimental setup for uniaxial tensile testing with DIC. Close-up view shows spray-painted plaque strip for adequate speckle pattern tracking.

#### iii. Histological analysis

After fixation, strips were stepwise dehydrated (Leica TP1020, semi-enclosed bench-top tissue processor, Germany) and embedded in paraffin wax blocks. Samples were sectioned at 5 μm to obtain axial cross-sections using Feather C35 microtome blades and stained with haematoxylin & eosin (H&E). Intraplaque neovascularisation was evaluated using CD31 markers for immunohistochemistry. Deparaffinised sections were processed for antigen retrieval using a pressure cooker. The sections were incubated with the primary antibody against CD31 (mouse monoclonal, 1:50 dilution, Anti-CD31 antibody [JC/70A]; Abcam) overnight at 4°C. Sections were subsequently treated with the secondary antibody for 1 hour at room temperature (EnVision+/HRP, Mouse, HRP, K4001, Agilent). Finally, the liquid DAB chromogen (Liquid DAB+, 2-component system, K3468, Agilent) was applied for 6–8 minutes. Brightfield imaging on an Aperio CS2 microscope using ImageScope software V12.3 (Leica Biosystems Imaging, Inc., Vista, California) was then performed on the stained sections. The workflow is illustrated in Figure 5.

**Figure 5:**
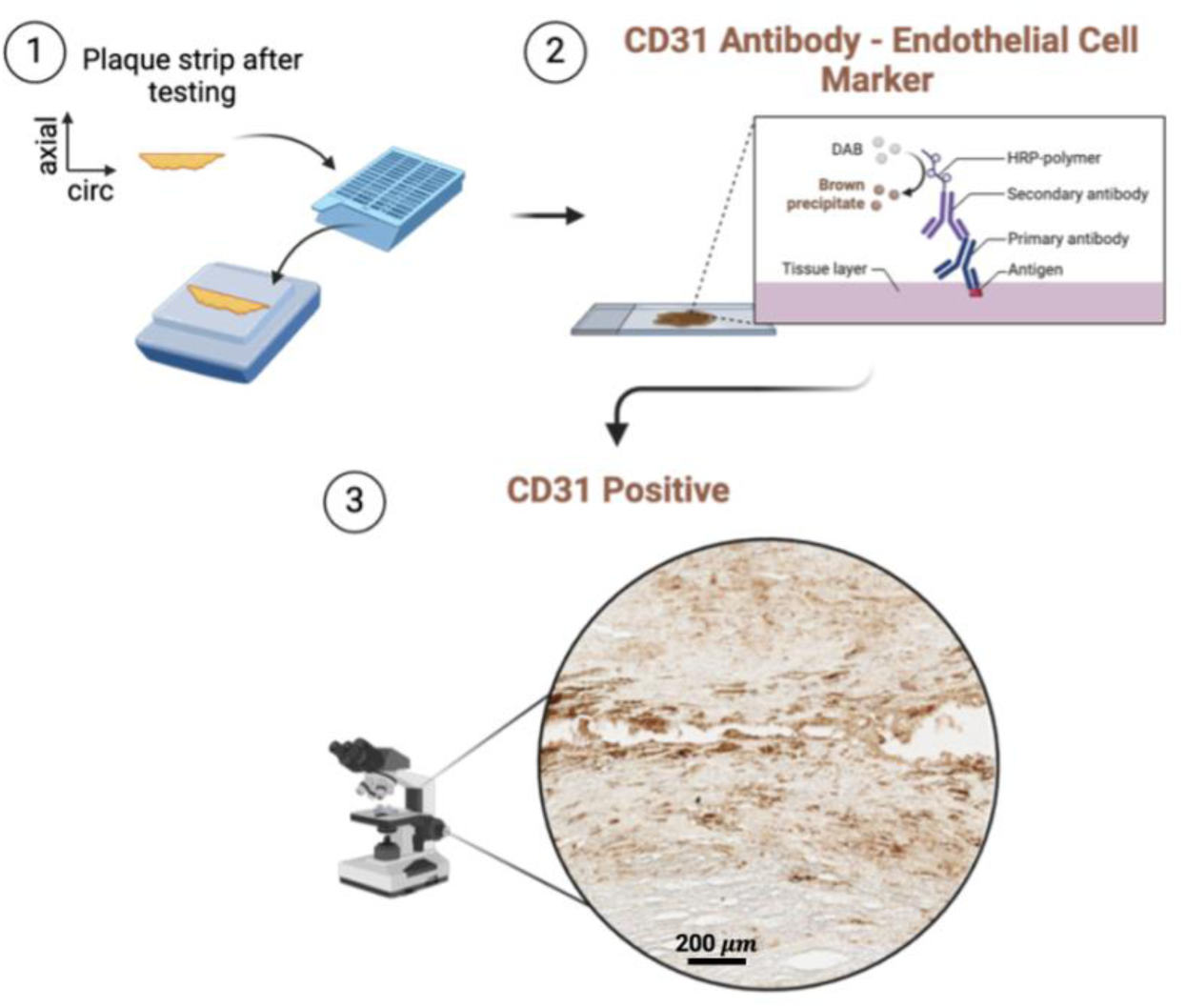
I*m*munohistochemistry *workflow to visualise neovessels in strips*.

#### iv. Neovascularisation quantification

Histology quantification was performed using QuPath (v 0.5.1). Digital whole immunohistochemistry slides were opened in QuPath, where the following procedure was followed to quantify the neovessel percentage per stained area. An automated thresholding approach was used to analyse positively stained plaque sections. This analysis employed an automated thresholding tool developed on <u>GitHub</u>, using the triangle method to identify neovessels. Consistent tissue regions were delineated using the Wand tool in QuPath, avoiding any folded areas or regions with background staining. The features of the thresholding algorithm were as follows: moderate resolution (3.96 μm/px) with DAB channel selected. Two sections per strip were analysed, with tissue regions delineated to exclude folds. Neovessel percentage was based on the stained area and was averaged across both slices to provide a measure of neovascularisation.

##### Mechanical data

Engineering stress was calculated using the cross-sectional area of each specimen. The force-displacement curves obtained after preconditioning, were analysed to determine the stress-strain behaviour of the samples. Failure was defined as the point at which a 5% decrease in stress between two consecutive points was first observed^23^. At this point, ultimate tensile (UT) stress and UT strain values were recorded. The final elastic modulus was calculated by fitting a line between the UT stress and 70% of the UT stress, while the initial elastic modulus was defined as the slope of the curve between zero and 5% strain^24^. DIC analysis was conducted using Istra4D software (x64 V4.4.6.534) with the following parameters: facet size of 69, 3D residuum of 10, grid spacing of 15 pixels, and a low outlier tolerance. The reference frame for all analyses was the final frame of the preconditioning cycles, just before extension to failure began. Engineering strain was assessed using DIC in two ways: (1) the average strain across the gauge length on the tissue surface, referred to as DIC strain (DIC-G), and (2) the average strain locally at the failure site, referred to as DIC local failure strain (DIC-L).

#### v. K-means clustering and decision tree classifier

A k-means clustering algorithm was implemented to classify strips based on UT stress and UT strain. These parameters were chosen due to their direct relevance to mechanical failure and clear interpretability from stress-strain curves. Evaluation of the impact of adding other mechanical features for k-means clustering are detailed in the supplementary data. UT stress and UT strain alone provided the best clustering performance, see Supplementary Figure 1, and as a result were used as the final k-means clustering approach.

The optimal number of clusters k was determined by evaluating multiple clustering validity indices, including the elbow method, silhouette analysis, Davies-Bouldin index, gap statistic, and Calinski-Harabasz index (Supplementary Figure 2). These metrics were computed for k values ranging from 2 to 4 to assess clustering robustness. Based on these analyses, k=2 was selected as the most appropriate cluster number, balancing intra-cluster compactness and inter-cluster separation.

Neovascularisation percentages for each strip, quantified from histological analysis described above, along with their cluster belonging were then extracted. To validate neovascularisation as a discriminative feature between the two clusters, a decision tree classifier was trained using neovascularisation percentages as the predictor variable and the k-means cluster labels as the response. The classification accuracy was calculated as:

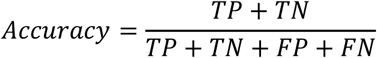

where TP refers to true positives, TN to true negatives, FP to false positives, and FN to false negatives. True positives and true negatives indicate correctly classified samples within their respective clusters, while false positives and false negatives represent misclassified samples. A receiver operating curve (ROC) curve was plotted, and the area under the curve (AUC) was calculated to quantify the discriminative ability of the classifier.

### c. Histological classification of plaque cross-sections

To complement the mechanical characterisation of plaque strips, additional plaque samples were classified according to the Stary et al. framework, which categorises atherosclerotic lesions based on their structural and compositional features^46^. Plaque samples (n=5) were cut into 2 mm thick rings using a 3D-printed blade holder and fixed in 10% formalin for histological processing as described in section(b)iii. Plaque samples were classified using the histological information gathered through H&E-based morphological features. This morphological assessment provided the foundation for categorising each plaque sample within Stary’s classification framework.

### d. Statistical analysis

Statistical analyses were performed using GraphPad Prism (Version 10). All data were tested for normality using the D’Agostino-Pearson test and for equality of variances using Brown-Forsythe ANOVA. If normality was not met, non-parametric tests were applied. Pearson’s correlation coefficient *r* was used to assess relationships between mechanical properties and neovascularisation, with correlation strengths defined as follows: weak (r < 0.3), moderate (0.3 < r < 0.7), and strong (r > 0.7). Outliers were identified using the ROUT method (maximum false discovery rate = 1%, Q = 1%). Any identified outliers were excluded from statistical analysis but are displayed in supplementary data. Results are presented as mean ± standard deviation (SD) for t-tests, median with interquartile range (IQR) for Mann-Whitney tests, and median with 95% confidence intervals (CI) in figures where applicable.

## 3. Results

Across the 25 strips, variability in mechanical properties was observed, as illustrated in Figure 6(A-D). This variability is further evidenced by the different responses to load observed in the stress-strain curves, see in Figure 6(E). The average UT stress and strain for the samples was 0.47 ± 0.46 MPa and 32.6 ± 18.5% strain, respectively. The mean initial modulus was 0.54 ± 0.63 MPa, while the mean final elastic modulus across all samples was 2.30 ± 2.28 MPa.

**Figure 6:**
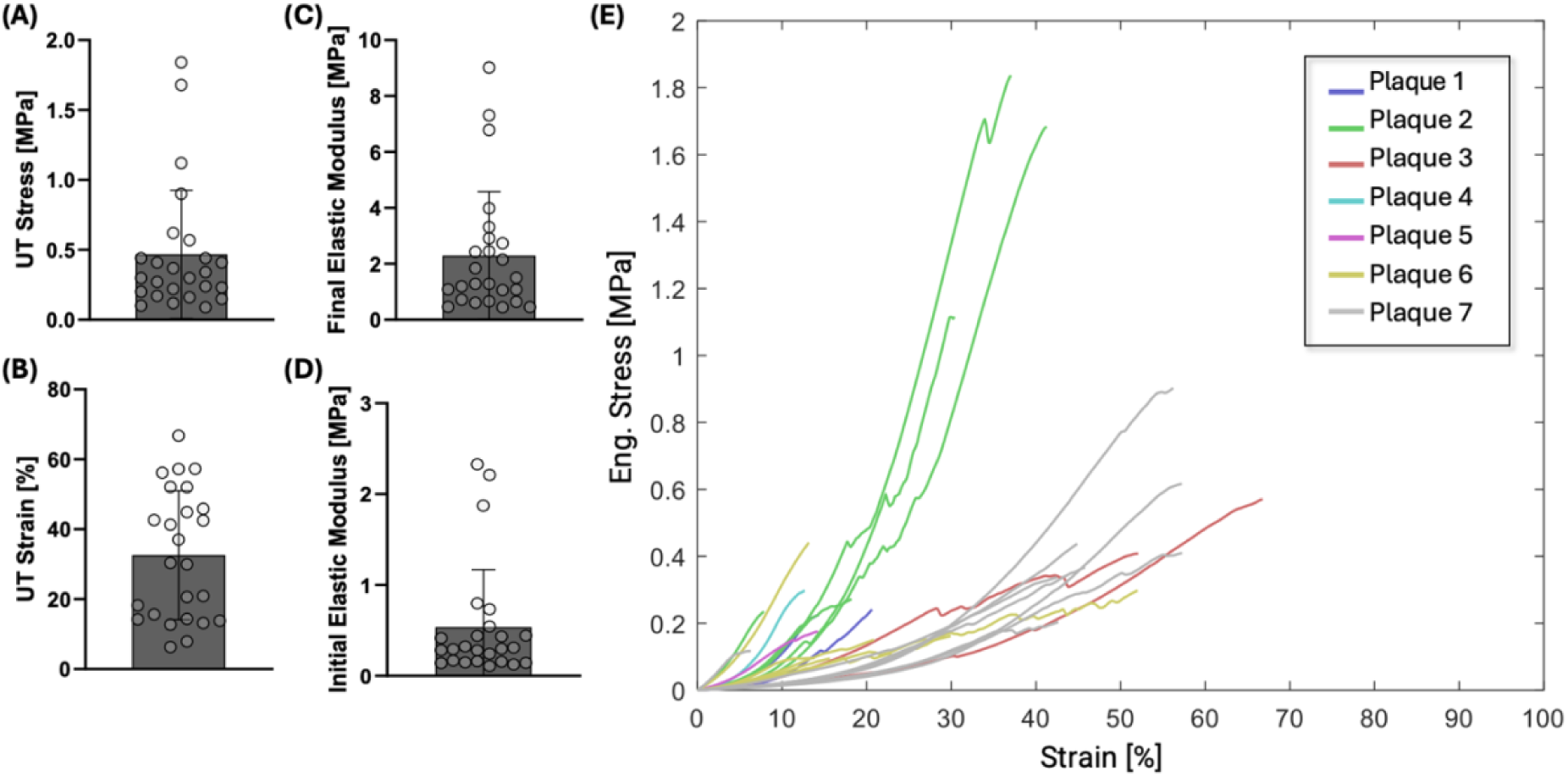
Mechanical properties of carotid atherosclerotic plaque strips (n=25). (A) UT Stress [MPa], (B) UT Strain [%], (C) final elastic modulus [MPa], (D) initial elastic modulus [MPa] and (E) engineering stress-strain curves for all strips colour coded by their respective plaque provenance. Note that SD exceeds the lower limit for the initial elastic modulus, resulting in clipped error bars at zero.

DIC allowed for visualisation of the progression of strain from the initial state to the point of rupture, with localised strain significantly higher at failure sites compared to the mean strain across the tissue surface, see Figure 7. This is also shown quantitatively by comparing localised strains at failure points, DIC-L, to DIC-G strain, where DIC-L was significantly higher than DIC-G. Additionally, the grip-to-grip strain from the testing machine significantly overestimated the DIC-G strain on the tissue surface.

**Figure 7:**
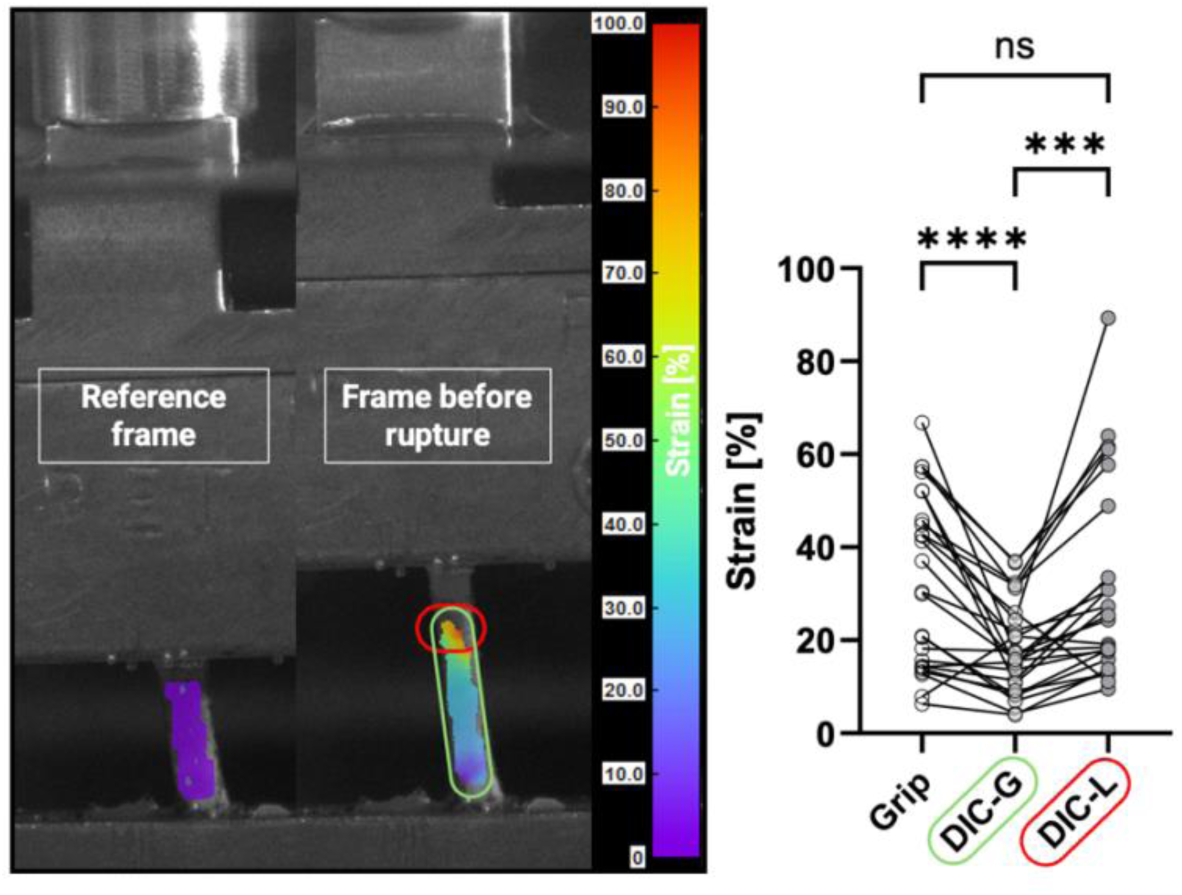
Strain maps at the reference frame and the frame before rupture, with DIC highlighting localised strain concentrations at failure (circled in red) and strain distribution across the surface (circled in green). Grip-to-grip strain (Grip), mean DIC strain across the tissue surface (DIC-G), and localised DIC strain at failure sites (DIC-L) are compared, demonstrating significant differences between these measurements. Statistical significance was assessed using repeated measures one-way ANOVA with Tukey’s post hoc multiple comparisons; ***p < 0.001, ****p < 0.0001.

Notably, strain maps revealed that rupture locations coincided with regions of maximum localised strain in n=20 out of the total n=25 tested strips, emphasising the role of localised strain concentration in mechanical failure. The relationship between UT stress and localised DIC-L strain at failure is further illustrated in Supplementary Figure 3.

Figure 8 shows the correlations between neovascularisation percentages and mechanical properties extracted from uniaxial testing. A weak negative correlation was observed between UT stress and neovascularisation (r = -0.2412). In contrast, UT strain showed a statistically significant moderate negative correlation (r = -0.4585, *p = 0.0278) with neovascularisation percentage. This indicates that strips with higher neovascularisation tend to fail at lower strains. The final elastic modulus exhibited no meaningful correlation with neovascularisation (r = -0.0376) and the initial elastic modulus displayed a negligible positive correlation (r = 0.0468), suggesting no significant interaction in either case. Among all mechanical properties analysed, UT strain demonstrated the strongest relationship with neovascularisation. Additionally, the area corresponding to UT strain values below 20% has been shaded in red in Figure 8(B), representing a threshold for typical physiological circumferential strain in human carotid arteries. These findings suggest that plaques rich in neovessels may rupture under normal or even sub-physiological loading conditions.

**Figure 8:**
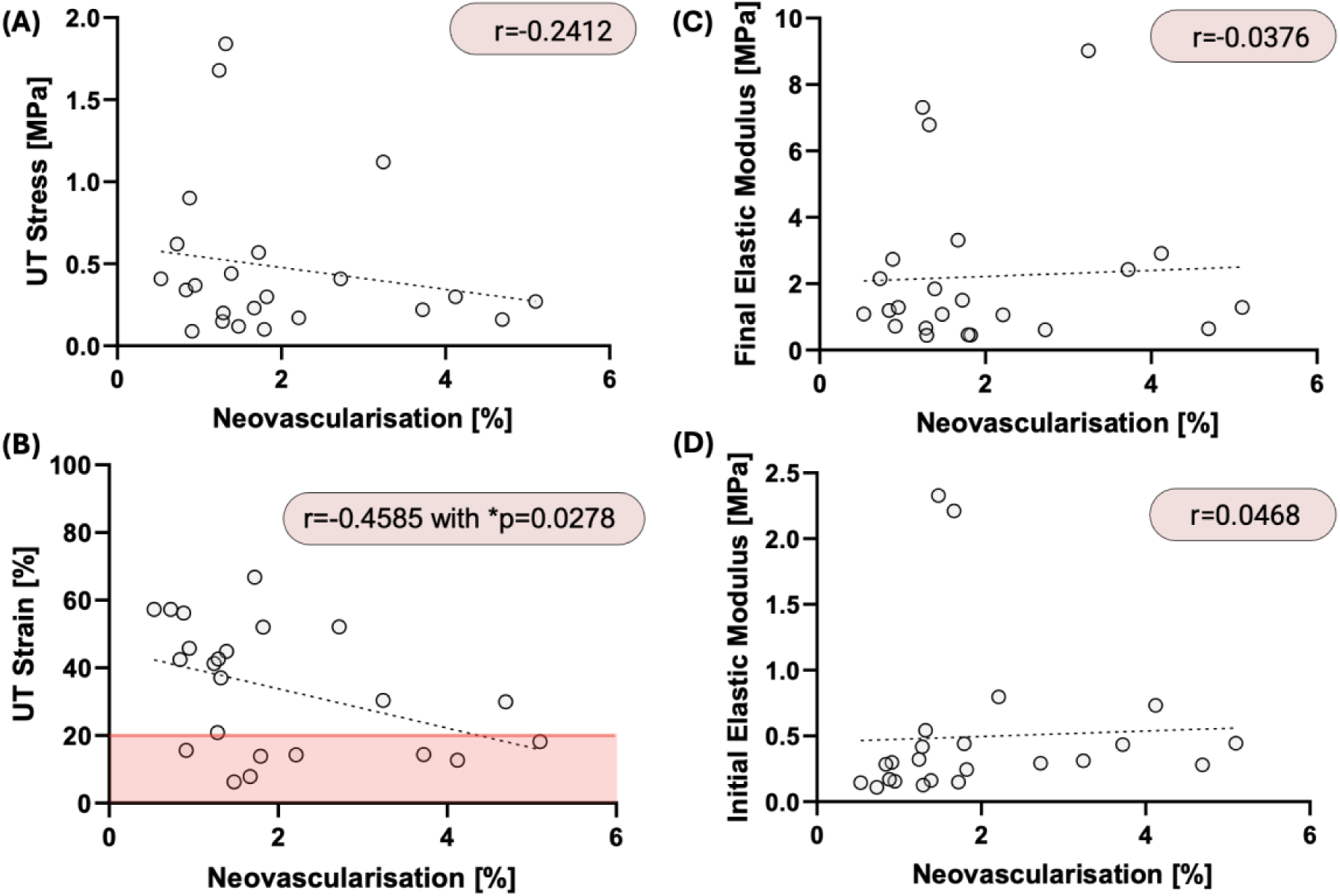
Scatter plots showing correlations between neovascularisation percentage and (A) UT stress, (B) UT strain, note that the shaded red region highlights the area below 20% UT strain, corresponding to the average physiological circumferential strain range typically experienced in human carotid arteries., (C) final elastic modulus, and (D) initial elastic modulus for n=23 strips. Dashed lines represent linear regression fits. Scatter plots showing correlations between neovascularisation percentage and (A) UT stress, (B) UT strain, (C) final elastic modulus, and (D) initial elastic modulus for n=23 strips. Dashed lines represent linear regression fits. Correlation coefficients ($r$) are provided, with significant correlations denoted by an asterisk (*p < 0.05). Note that *neovascularisation was quantified for each plaque, and after performing outlier analysis, one data point was excluded. Additionally, due to histological processing, some sections were not analysable and therefore excluded from the analysis, leaving a final sample size of n = 23*.

Additionally, a moderate negative correlation (r = -0.6420, ***p < 0.001) was found between UT strain and the initial elastic modulus, see Figure 9. This trend suggests that strips with higher initial elastic modulus tend to fail at lower strains.

**Figure 9:**
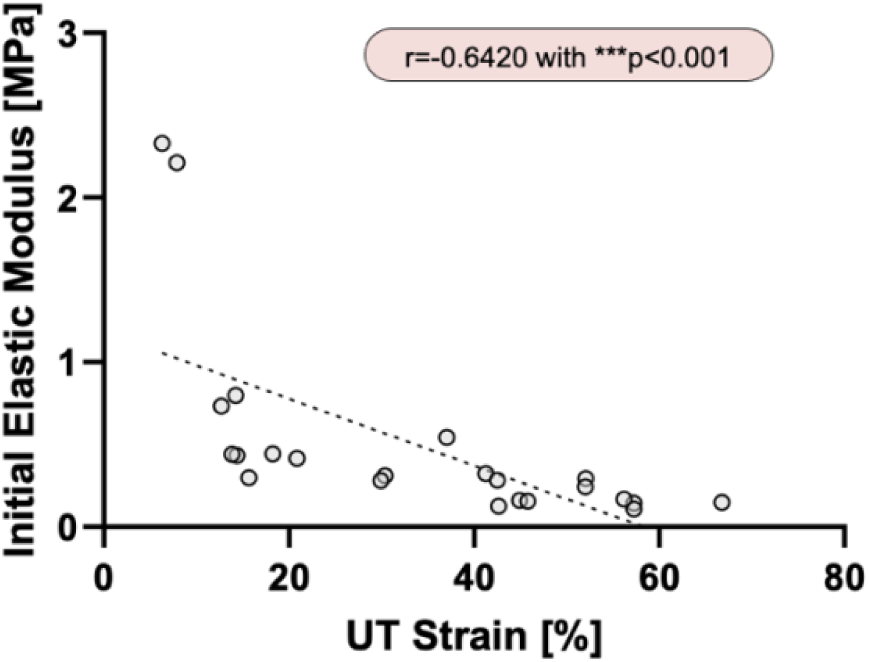
Scatter plot showing correlation between UT strain and initial elastic modulus of n=23 strips.

Figure 10 illustrates the relationship between UT strain and UT stress, with bubble sizes representing the percentage of neovascularisation. The plot highlights a distribution pattern where plaque strips exhibiting lower UT strain tend to have higher neovascularisation percentages, as shown by the cluster of larger bubbles below 20% UT strain. This observation suggests that increased neovascularisation is associated with reduced strain at failure resonating with findings seen in Figure 8(B).

**Figure 10:**
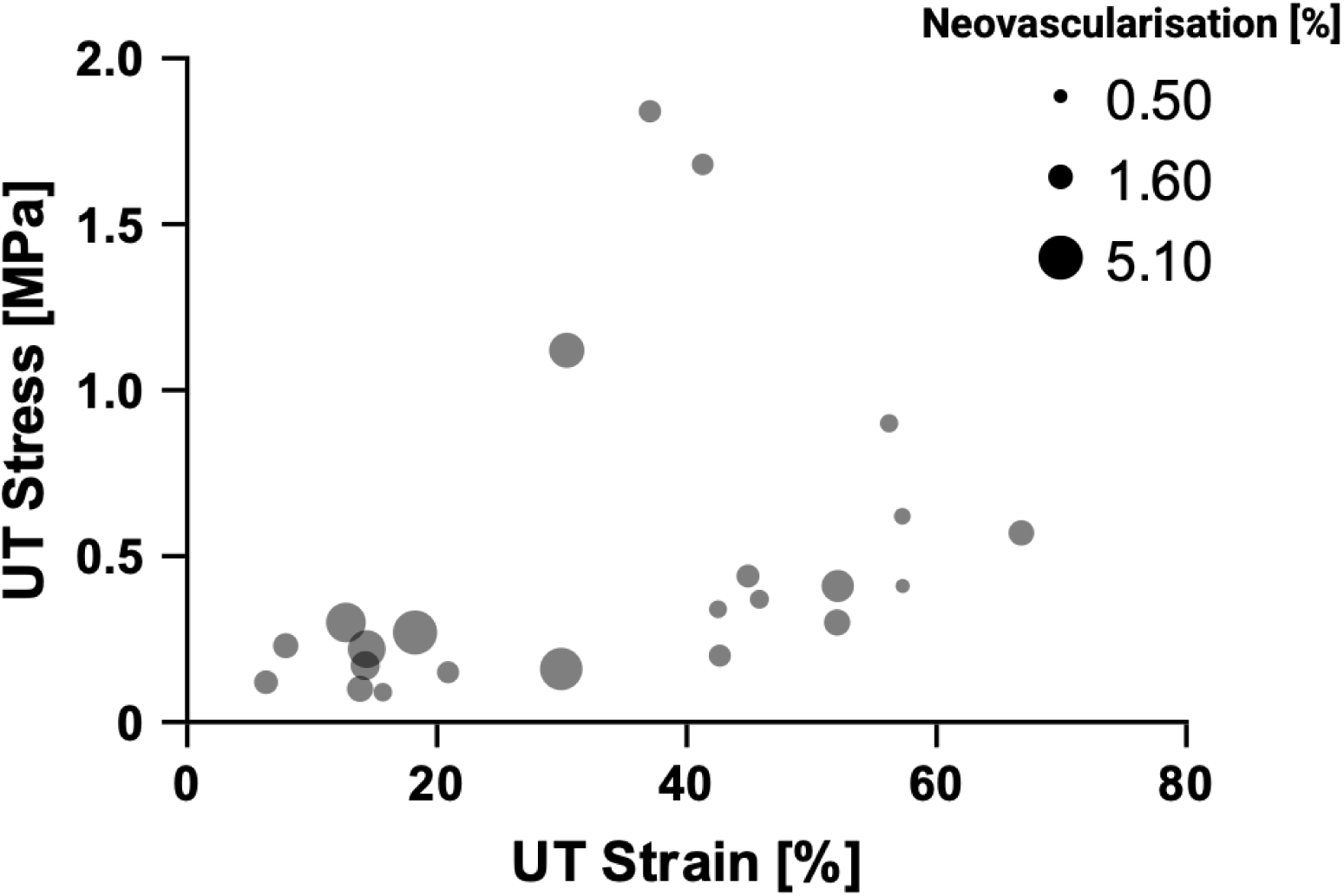
Bubble plot showing the relationship between UT strain and UT stress, with bubble sizes representing the quantification of neovascularisation percentages. The red ellipse highlights a cluster of plaques with low UT strain where larger bubbles indicate higher neovascularisation percentages.

To gain deeper insights, k-means clustering was applied (k=2) based on UT stress and UT strain. Figure 11 provides an overview of the clustering results. Panel (A) shows the stress-strain curves for each sample coloured by cluster (cluster 1: red, cluster 2: blue) with the cluster centroids marked by black crosses. Panel (B) further visualises the two clusters with each a specific plaque strip colour-coded by plaque sample. The clustering revealed two distinct behaviours within the dataset. Cluster 1 primarily consists of samples with lower UT strain values, highlighting a group of plaques that fail earlier under tensile loading. In contrast, cluster 2 includes samples capable of withstanding higher strains before failure. Interestingly, some strips from individual plaques predominantly belonged to a single cluster. Strips from plaque 6 predominantly belonged to cluster 1, reflecting its distinct mechanical properties characterised by withstanding lower UT strain. In contrast, strips from plaque 7 were largely grouped in cluster 2 – indicative of the ability to withstand higher strain before failing. However, overlap was observed for strips from plaques 2 with some strips appearing in both clusters, suggesting that these plaques exhibit mixed mechanical characteristics.

**Figure 11:**
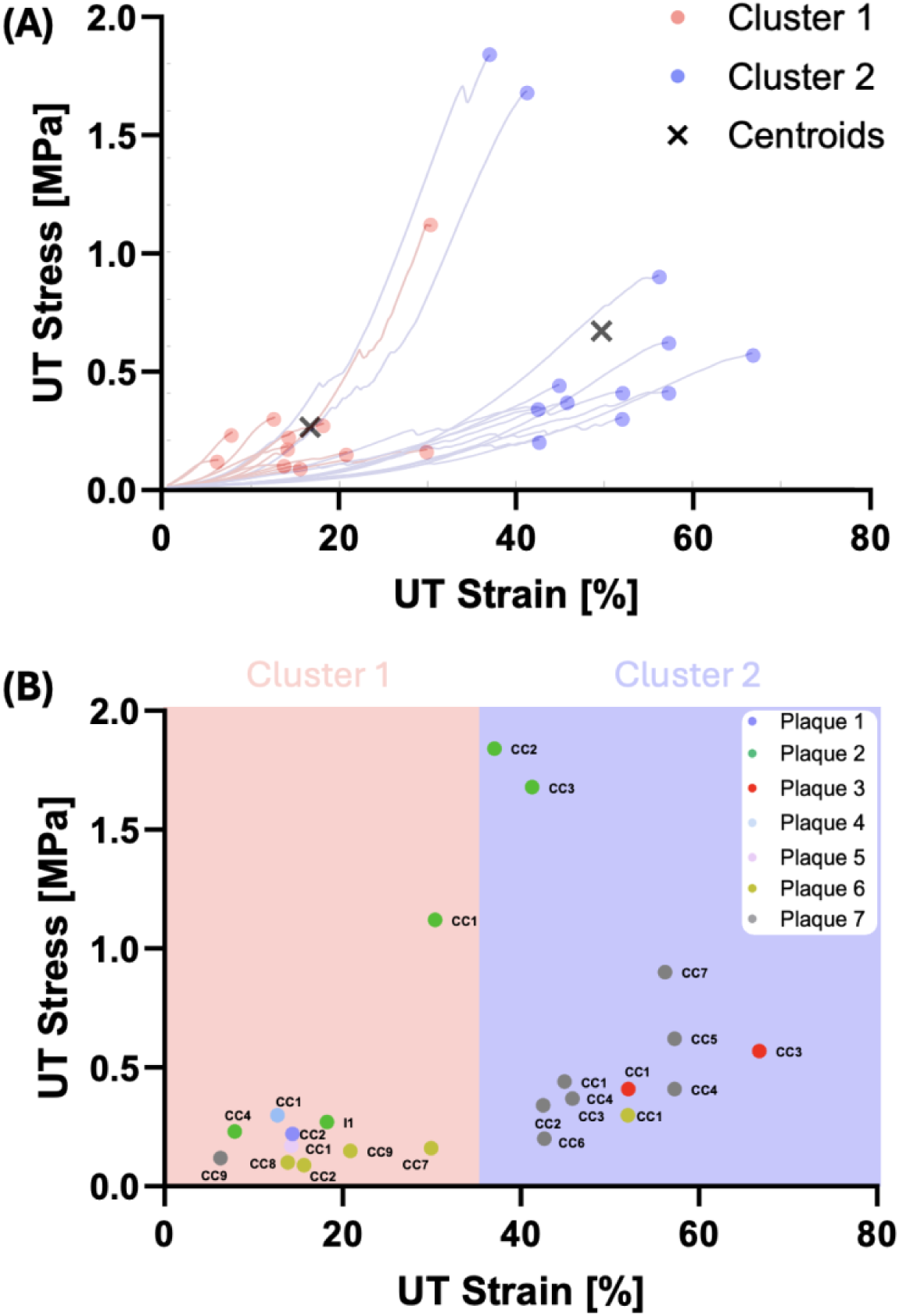
(A) Stress-strain curves for strips clustered into two groups using k-means clustering (k=2, cluster 1: red, cluster 2: blue). Cluster centroids are marked with black crosses. (B) Scatter plot of UT strain versus UT stress, showing the distribution of samples across the two clusters. Each point is colour-coded by plaque identifier, highlighting the distinct mechanical behaviours of the two groups. CC: common carotid, I: internal.

To further characterise the strips within each cluster, Figure 12 integrates the k-means clustering results with histological observations. Representative histological sections, including H&E and CD31 staining, are shown for strips closest to each cluster centroid. The H&E staining highlights areas of heterogeneity within one strip from cluster 1, specifically at the rupture region. CD31 staining reveals increased neovascularisation, particularly in this thickened, disorganised zone. In contrast, a representative strip in cluster 2, displays less complex and more homogeneous tissue structures. The H&E staining reveals more organised tissue layers and CD31 staining further demonstrates reduced neovascularisation. Red rectangles in the figure highlight magnified zones for a closer examination of tissue organisation where rupture occurred showing no presence of neovascularisation for the strip in cluster 2. These observations are further shown in supplementary Figure 4 and supplementary Figure 5.

**Figure 12:**
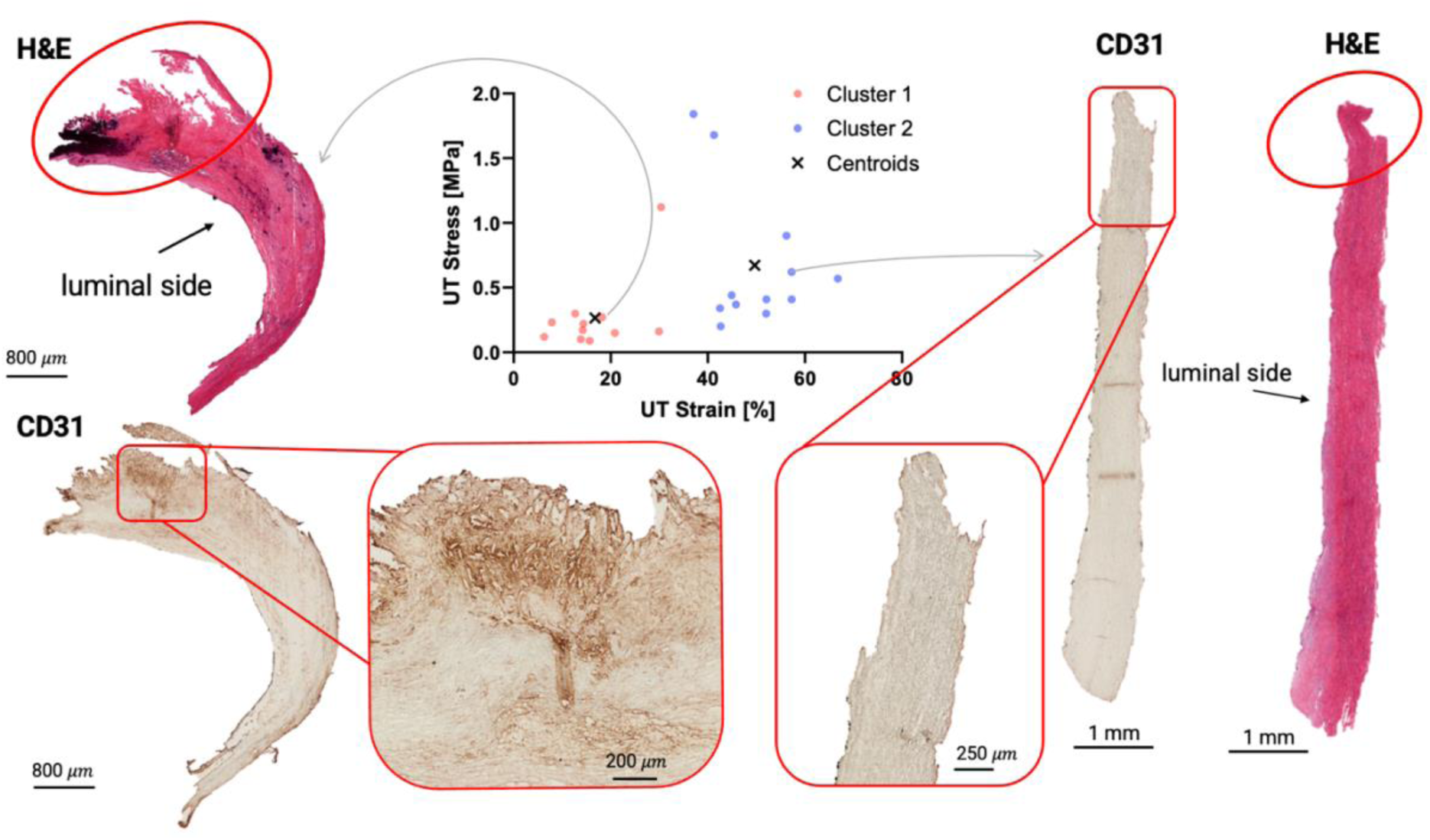
Combined k-means clustering of UT stress versus UT strain with representative histological sections (H&E and CD31 staining) for strips closest to each cluster centroids. Scatter plot shows the two clusters (cluster 1: red, cluster 2: blue) with centroids marked as black crosses. Red rectangles indicate magnified zones for detailed visualisation of tissue organisation and neovascularisation.

Figure 13 compares neovascularisation, DIC-L, the final elastic modulus, and the initial elastic modulus between clusters. Neovascularisation percentages were significantly higher (**p < 0.01) in cluster 1 (Median = 2.21%, IQR = 2.64) compared to cluster 2 (Median = 1.27%, IQR = 0.79), see Figure 12(A). For DIC-L, a significant difference (**p < 0.01) was found, with cluster 2 (43.92% ± 23.34) exhibiting significantly higher values than cluster 1 (20.60% ± 7.62%), see Figure 12(B). This indicates that strips in cluster 2 experience greater strain at rupture compared to those in cluster 1. The final elastic modulus showed no significant difference between clusters (cluster 1: 1.46 ± 1.04 MPa, cluster 2: 1.31 ± 1.31 MPa), see Figure 12(C). The initial elastic modulus was significantly higher (**p < 0.01) in cluster 1 (Median = 0.43 MPa, IQR = 0.28) compared to cluster 2 (Median = 0.17 MPa, IQR = 0.15), see Figure 12(D). This suggests that strips in cluster 1 exhibit greater initial stiffness but fail at lower strains compared to cluster 2.

**Figure 13:**
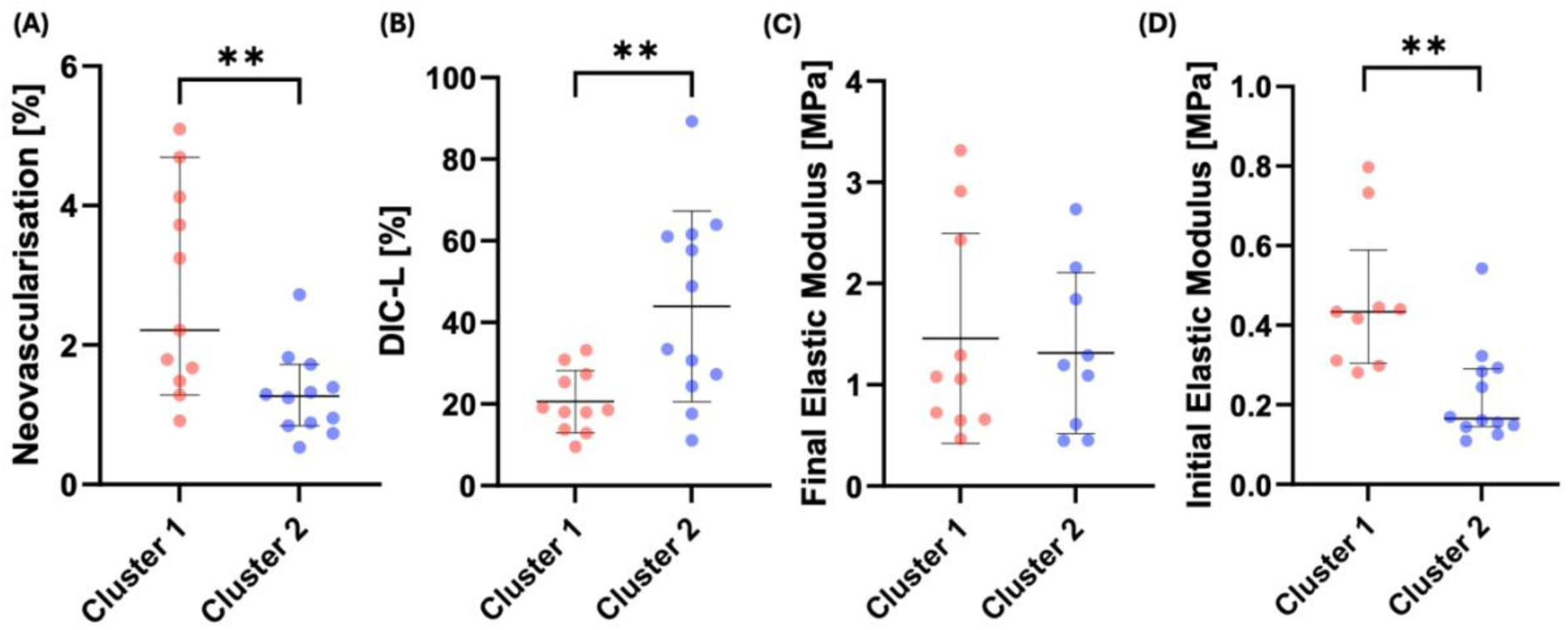
Comparison of mechanical and structural properties between cluster 1 (red) and cluster 2 (blue). (A) Neovascularisation percentages, analysed using the Mann-Whitney test **p < 0.01), reported as median with 95% CI by black horizontal lines. (B) DIC-L, analysed using a t-test (**p < 0.01), reported as mean ± SD by black horizontal lines. (C) Final elastic modulus, analysed using a t-test (p = 0.74), reported as mean ± SD by black horizontal lines. (D) Initial elastic modulus, analysed using the Mann-Whitney test (**p < 0.01), reported as median with 95% CI by black horizontal lines. Outliers were observed in initial and final elastic modulus measurements for cluster 1. These outliers were not included in the statistical analysis but are presented in the supplementary data for completeness, see Supplementary Figure 6.

The decision tree classifier identified a neovascularisation threshold of 1.44% as the optimal cut-off for classifying the two clusters. This threshold yielded a classification accuracy of 78.26%, suggesting that neovascularisation alone provides a reasonable, though not perfect, discriminant metric for mechanical clustering. To further assess the classification performance of neovascularisation as a discriminant metric between clusters, a ROC curve was generated (Figure 14). The ROC curve illustrates the trade-off between sensitivity (true positive rate) and specificity (false positive rate) in classifying plaques into the two clusters. The AUC was found to be 0.78, indicating that the decision tree model performed significantly better than random guessing.

**Figure 14:**
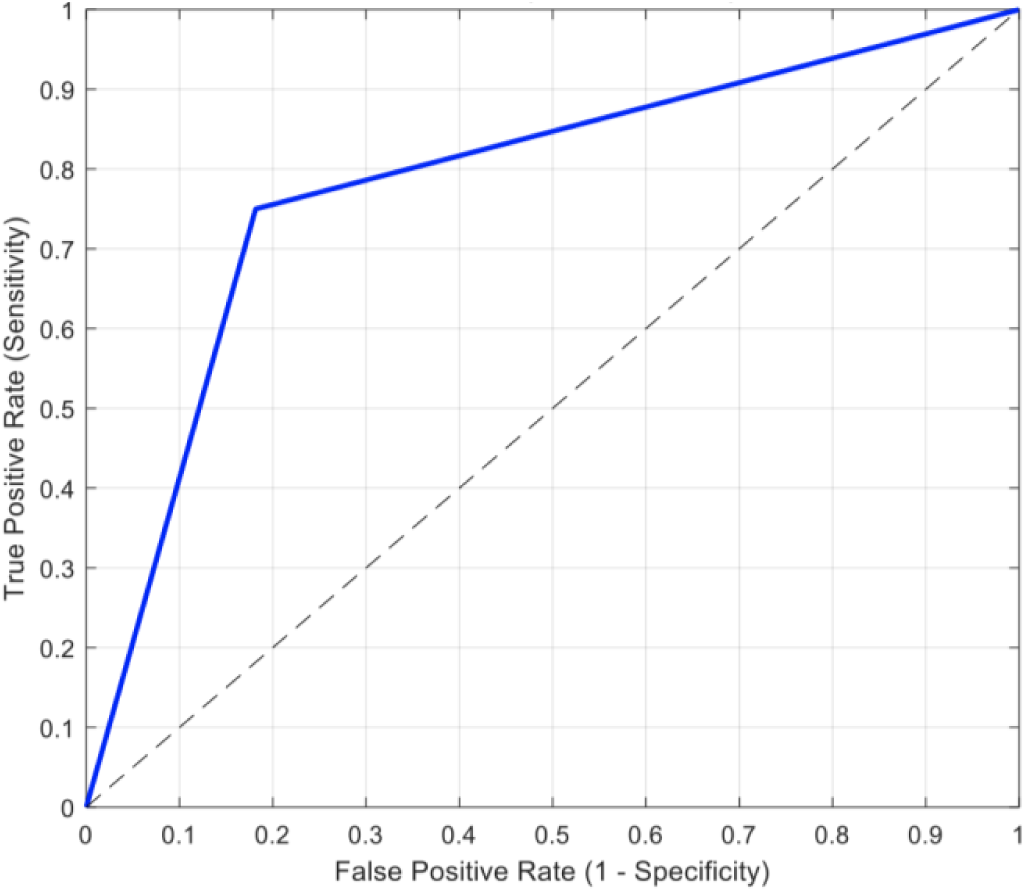
Decision tree classification of plaques based on neovascularisation percentage. The optimal threshold of 1.44% yielded a classification accuracy of 78.26%.

## 4. Discussion

This study provides novel insights into the relationship between neovascularisation and the mechanical properties of carotid atherosclerotic plaques. Neovascularisation has long been recognised as a key feature of atherosclerotic plaque instability. Virmani et al. and other studies have emphasised neovascularisation’s critical role in advancing plaque vulnerability by facilitating microvessel formation and intraplaque haemorrhage^10,17,21^. These highlight how neovascularisation has remained a focal point in understanding plaque instability over the years, emphasising its critical potential in identifying and addressing vulnerable plaques.

Multimodal US techniques have significantly advanced the assessment of carotid atherosclerotic plaques. Among these techniques, CEUS has emerged as a powerful tool for visualising intraplaque neovascularisation^16^. The ability of CEUS to capture and quantify intraplaque neovascularisation (IPN) highlights its utility in both research and clinical settings, where it can complement other imaging modalities like SWE^18^. Feinstein et al. highlighted that CEUS enables real-time visualisation of plaque neovascularisation and the adventitial vasa vasorum, proposing its use as a non-invasive tool to enhance the assessment of carotid plaque and improve clinical management ^19^. Huang et al. found that neovascularisation originates predominantly from the adventitial vasa vasorum linking vascular remodelling to plaque instability^25^. This joins Moulton’s review which examines the role of plaque angiogenesis in atherosclerosis, highlighting that neovascularisation within atherosclerotic plaques primarily arises from the adventitial vasa vasorum and extends into the intima^26^. Additionally, studies have shown the high correlation between histology-based neovascularisation and CEUS^14,20^.

Many studies in the literature have tackled the question inquiring the role that neovascularisation plays in atherosclerosis. Shah et al. emphasised the critical association between IPN and plaque instability, with a particular focus on the impact of radiotherapy (RT)^21^. Their findings demonstrated that plaques in patients with prior RT exhibited significantly increased IPN compared to non-RT plaques. Interestingly, RT is strongly associated with a heightened risk of ischaemic stroke, potentially due to the presence of neovascularisation and its contribution to plaque instability. However, few studies have explored the connection between plaque mechanical behaviour and neovascularisation, leaving this area of research still relatively scarce^27–30^.

Zhang et al. showed the predictive power of SWE and CEUS for stroke^27^. Wang et al. found that plaques with lower echogenicity and lower Young’s modulus showed increased neovascularisation^28^. Tian et al. used CEUS and found increased neovascularisation in soft plaques^30^. On the other hand, Zhang et al. found a negative correlation between neovascularisation and elasticity, where increased neovascularisation was indicative of lower elasticity^29^. This aligns with our findings, where strips with higher neovascularisation were associated with greater initial elastic modulus (Figure 8(D)). The variability in the mechanical response of carotid plaques observed in this study is consistent with previously reported findings^5,6,8,31–34^.

Loree et al. reported tangential moduli of 0.927 ± 0.468 MPa for cellular, 2.312 ± 2.180 MPa for hypocellular, and 1.466 ± 1.284 MPa for calcified plaques under a tensile stress of 0.025 MPa^31^. These values overlap with our final elastic modulus (2.30 ± 2.28 MPa). Similarly, Tornifoglio et al. found a mean modulus of 1.26 ± 0.6 MPa, consistent with our results^8^.Our UT stress values (0.47 ± 0.46 MPa) align with those reported by Lawlor et al. (0.131–0.779 MPa), Johnston et al. (0.31 ± 0.18 and 0.87 ± 0.63 MPa), Guvenir et al. (0.12–0.36 MPa), and Tornifoglio et al. (0.293 ± 0.2 MPa)^5,6,8,35^. For UT strain, our mean (32.6 ± 18.5%) falls within the range reported by Lawlor et al. (Green strains: 0.299–0.588), Johnston et al. (0.13 ± 0.04 and 0.09 ± 0.04), Guvenir et al. (stretch ratios: 1.16–1.64), and Tornifoglio (38.3 ± 19%)^5,6,8,35^.

Another interesting observation relates to the rupture initiation regions reported by Guvenir et al. which exhibited significantly higher stretch ratios (1.26 [1.15–1.40]) compared to the tissue’s average stretch ratio (1.11 [1.10–1.16])^6^. This aligns with our findings that strain at rupture (DIC-L) was significantly higher than strain across the tissue surface (DIC-G). A similar observation was made in the study by Tornifoglio et al., further supporting the idea that strain could serve as an indicator of plaque vulnerability^36–38^.

In a recent study, Crielaard et al. performed uniaxial tensile testing on tissue-engineered fibrous cap analogues and demonstrated that rupture initiates and propagates through regions of high strain, mirroring patterns observed in human plaques^39^. This observation is further supported by our findings, where DIC revealed that rupture locations consistently coincided with regions of maximum localised strain in n=20 out of the total n=25 tested strips. The coincidence of rupture location and high strain concentration highlights the importance of incorporating localised strain metrics when assessing plaque vulnerability, as these regions may represent focal points for mechanical instability and potential rupture.

Adding to this, correlation analysis in this study revealed that UT strain demonstrated the strongest association with neovascularisation with statistically significant moderate negative correlation (r = -0.4585, *p = 0.0278). This moderate correlation between UT strain and neovascularisation underscores its role on plaque mechanical vulnerability: highlighting the critical importance of integrating mechanical metrics with microstructural information. UT strain was also found to be significantly correlated to the initial elastic modulus (r = -0.6420, ***p < 0.001) where lower UT strain is associated with higher initial elastic modulus.

Furthermore, the scatter plot reveals a notable pattern where larger bubbles, representing higher neovascularisation percentages, seem predominantly associated with lower UT strain values (Figure 10). This observation suggests that plaques with greater neovascularisation may rupture at lower strain, further reinforcing the interplay between microstructural features and mechanical properties in plaque vulnerability. A closer examination of the scatter plot through that k-means clustering further highlights this behaviour within the dataset. Cluster 1, characterised by lower UT strain, was associated with higher neovascularisation percentages, and demonstrated significantly higher initial elastic modulus (Figure 13(A, D)). In contrast, strips in cluster 2 displayed higher UT strain, as well as lower initial elastic modulus (Figure 13(A, D)).

Such observations support the hypothesis that neovascularisation, through its association with immature and leaky microvessels contributes to overall plaque instability^10,17^. Teng et al. investigated the mechanical environment surrounding neovessels and found that neovessels experienced the highest levels of stress and stretch during the cardiac cycle through finite element analysis and histological analyses^23^. These mechanical conditions around neovessels are potentially driving the progression of IPH. This highlights the critical role of neovascularisation in understanding and managing plaque vulnerability. Supporting this, Saba et al. emphasised the importance of neovascularisation as a factor in the RADS scoring system, further linking it to an increased risk of plaque rupture^4^. Li et al. used numerical simulations based on CEUS images to evaluate stress and strain distribution within carotid atherosclerotic plaques^40^. Their findings indicate that neovessels within plaques are particularly prone to deformation, which can lead to rupture. Altogether, these findings highlight the power of integrating plaque mechanical and microstructural characteristics to enhance risk assessment.

In this study specifically, combining neovascularisation and mechanical properties highlighted the potential to provide deeper insights into plaque vulnerability. The findings in this study suggest that neovascularisation may contribute to mechanical instability. Neovascularisation was moderately negatively correlated with UT strain which was reinforced by objective k-means clustering which revealed that strips with lower strain at failure exhibited significantly higher neovascularisation and greater initial elastic modulus. To further assess the discriminative power of neovascularisation in classifying mechanical behaviour, a decision tree classifier was implemented using neovascularisation percentage as the sole predictor variable. The classification results revealed an optimal threshold of 1.44% neovascularisation for distinguishing between the two k-means clusters, achieving an overall classification accuracy of 78.26%. This suggests that neovascularisation alone provides a reasonable, though not absolute, metric for stratifying plaques based on their mechanical properties. Additionally, the ROC curve analysis demonstrated an AUC of 0.784, further supporting the ability of neovascularisation to differentiate between plaques with lower and higher UT strain at failure. Together, these findings demonstrate that neovascularisation may contribute to mechanical instability, supporting its role as a potential marker for plaque mechanical integrity.

This study is, however, subject to some limitations that warrant consideration. First, uniaxial tensile testing, which was conducted on *ex vivo* samples, does not fully replicate the physiological *in vivo* conditions. However, the strips were tested axially to replicate physiological loading in the circumferential direction, making the findings still relevant to understanding plaque behaviour under loading. Secondly, the quantification of neovascularisation relied on histological analysis, which provides only a partial measurement of its spatial variability through the whole plaque. Specifically, the analysis was based on two histological slices per plaque, with neovascularisation averaged across these sections. This approach may not fully capture the heterogeneous distribution of neovascularisation throughout the entire plaque. Nonetheless, the moderate but significant correlation observed between UT strain and neovascularisation from immunohistochemistry offers promising insights into the relationship between neovascularisation and plaque instability.

Future studies should prioritise *in vivo* evaluation of neovascularisation using readily available techniques like CEUS, which can provide a more comprehensive view of neovascularisation distribution and its association with mechanical properties. Such imaging methods could enhance our understanding of plaque vulnerability and improve clinical risk stratification. Additionally, ongoing advancements in US imaging, such as SMI, could enable non-contrast quantification of neovascularisation, making these techniques more accessible for routine clinical use^41–44^.

While this study focused on individual strips, the overlap of plaques between clusters suggests that strip-level analysis may not fully capture the mechanical behaviour of entire plaques. Some plaques predominantly contributed to one cluster, while others exhibited characteristics spanning the two clusters. This variability highlights the importance of whole-plaque studies that integrate mechanical testing with imaging to better understand the spatial heterogeneity of plaque properties.

As mentioned, the relationship between plaque neovascularisation, progression, and symptoms remains complex and unresolved. For example, while inflammation has been shown to be more pronounced in symptomatic patients, neovascularisation does not consistently correlate with symptoms^45^. On the other hand, Nardi et al. found that symptomatic plaques were associated with IPH, which is closely linked with neovascularisation^13^.

### Plaque Classification and neovascularisation trends

Unlike the mechanically tested strips, the plaque samples (n=5) that were sectioned in cross-sections, allowed for a direct evaluation of plaque morphology and neovascularisation distribution across the lumen. Figure 15 presents histological sections of the plaques, stained with CD31 for neovascularisation (top row) and H&E for plaque morphology (bottom row).

**Figure 15:**
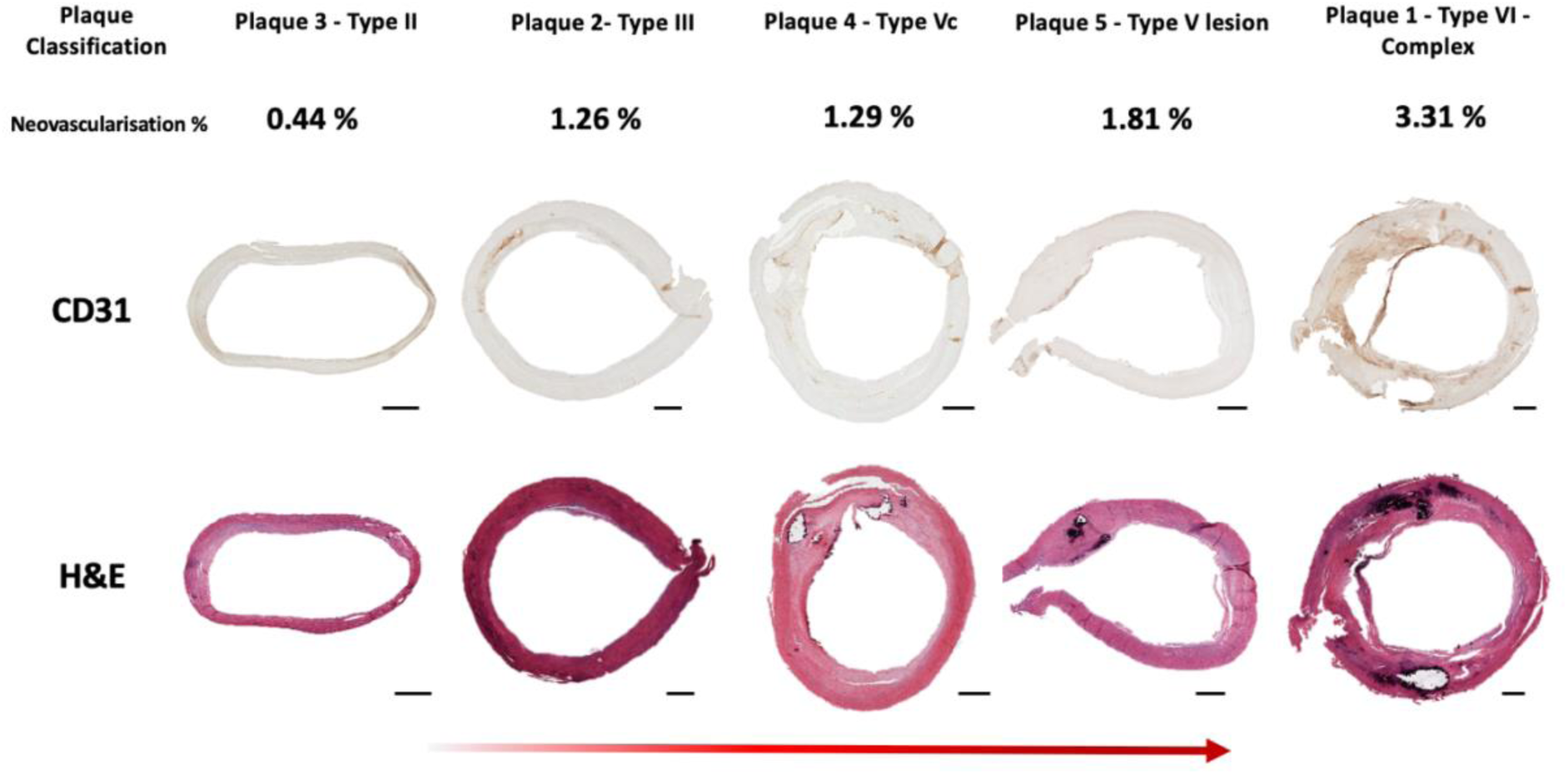
Histological sections of atherosclerotic plaques classified using the Stary framework\cite{stary1995definition}. The top row displays CD31-stained sections highlighting neovascularisation. The bottom row shows H&E-stained. Scale bars: 1 mm.

This classification highlights a trend of increasing neovascularisation with plaque complexity:

- ***Plaque 3 (Type II):*** Minimal structural complexity, sparse CD31-positive staining, low neovascularisation (0.44%).
- ***Plaque 2 (Type III):*** Slightly increased vascularisation (1.26%).
- ***Plaque 4 (Type Vc):*** Distinct lipid cores, thick fibrous cap, and calcifications with localised neovascularisation (1.29%).
- ***Plaque 5 (Type V):*** Presence of fibrous structures, necrotic core, and calcification, with neovascularisation in the inner lumen of the plaque (1.81%).
- ***Plaque 1 (Type VI - Complicated lesion):*** Most advanced plaque, with extensive CD31-positive staining and the highest neovascularisation percentage (3.31%).

This classification reinforces the progression of neovascularisation with plaque severity. Advanced plaques, such as Type V and Type VI lesions, exhibited greater neovascularisation, consistent with previous findings linking intraplaque neovascularisation to increased plaque vulnerability and intraplaque haemorrhage^10,17^.

To our knowledge, this is the first study to establish a link between neovascularisation and rupture mechanical metrics, specifically UT strain. We showed that plaques with higher neovascularisation have lower UT strain, indicating greater vulnerability. By integrating imaging-derived metrics with mechanical properties, clinicians could better stratify patients and optimise surgical interventions. This approach moves beyond the traditional reliance on stenosis as a primary risk metric, advocating for a multimodal assessment of plaque vulnerability. Given that neovascularisation is already measurable using clinical tools like CEUS, its incorporation into clinical workflows could facilitate improved risk assessment with minimal barriers to bridge the gap between the lab and the clinic.

## Acknowledgements

The authors would like to thank the Department of Vascular Surgery at the Galway Clinic for supplying the human tissue used in this study. Specifcally, Prof. Sherif Sultan and Dr Niamh Hynes from Galway Clinic. We also thank Kim van Gaalen for her assistance with the immunohistochemistry protocol. This work was conducted with the financial support of the Research Ireland Centre for Research Training in Digitally-Enhanced Reality (d-real) under Grant No. 18/CRT/6224. For the purpose of Open Access, the author has applied a CC BY public copyright licence to any Author Accepted Manuscript version arising from this submission.

